# Inhibitory-modulatory coupling generates persistent activity during working memory

**DOI:** 10.64898/2026.03.27.714602

**Authors:** Marissa L. Heintschel, Jingyue Xu, Dhruv Grover

**Affiliations:** Kavli Institute for Brain and Mind, University of California, San Diego, La Jolla, CA, USA; Shu Chien-Gene Lay Department of Bioengineering, University of California, San Diego, La Jolla, CA, USA

## Abstract

Working memory requires the stable maintenance of neural representations across temporal gaps, yet the circuit mechanisms that generate and stabilize persistent activity remain unsolved. Prevailing models emphasize recurrent excitation as the principal substrate of persistence, but how inhibitory and modulatory interactions shape the stability of temporal dynamics is unclear. Here, using trace conditioning in *Drosophila*, a working memory-dependent form of associative learning, we identify reciprocal inhibition as a circuit mechanism for sustaining persistent activity. In trace conditioning, a “trace” interval separates the conditioned and unconditioned stimuli, requiring maintenance of a neural representation across the trace interval, to support learning. Combining virtual-reality behavior, targeted neurogenetic perturbations, *in vivo* two-photon calcium imaging, and real-time neurotransmitter measurements, we uncover a reciprocal inhibitory microcircuit within the ellipsoid body that is selectively engaged during trace, but not delay (overlapping CS-US), conditioning. During the trace interval, ER2/4m neurons exhibit sustained activity, while reciprocally connected ER3/4d neurons show progressively strengthened suppression, forming a dynamically stabilized inhibitory loop. Disrupting GABA synthesis or reception within this circuit abolishes persistent activity and impairs trace learning, demonstrating the causal requirement for reciprocal inhibition in working memory maintenance. We further show that glutamatergic and nitric oxide signaling enhance inhibitory efficacy during the trace interval. *In vivo* neurotransmitter imaging reveals temporally structured dynamics in which glutamatergic signaling precedes and amplifies sustained GABAergic inhibition, consistent with modulatory stabilization of circuit persistence. Together, these findings identify reciprocal inhibition, reinforced by modulatory signaling, as a core circuit mechanism for dynamically stabilizing persistent neural representations. Our results challenge excitation-centric models of working memory and establish inhibitory-modulatory loops as a fundamental substrate for maintaining memory traces across time.

## Main

Persistent neural activity is widely considered the neural substrate of working memory^1,2^, yet how circuits maintain such activity across behaviorally relevant timescales remains debated. Classical models emphasize recurrent excitation as the primary driver of persistence, forming attractor-like states that can sustain activity without input^3^. Emerging theory, however, suggests that inhibition may play a central role in stabilizing these dynamics^4^, controlling temporal integration, and preventing runaway excitation^5^. Whether defined inhibitory microcircuits are necessary to generate and sustain persistent activity *in vivo* has not been directly tested.

Trace conditioning provides a powerful framework for isolating working memory mechanisms^6^. In contrast to delay conditioning, in which conditioned and unconditioned stimuli overlap, trace conditioning inserts a temporal gap between stimuli, requiring maintenance of a neural representation of the conditioned stimulus across the trace interval to support associative learning^7^. Because both paradigms engage shared sensory and associative pathways^8,9^, yet only trace conditioning requires sustained internal representations^10^, dissociating their circuit requirements enables identification of mechanisms specific to working memory maintenance^11,12^.

Computational studies predict that strengthening inhibitory coupling can slow network dynamics^13^, enhance stability^14^, and promote self-sustained activity even in the absence of ongoing input^15^. Models of trace conditioning similarly indicate that shifts in inhibitory balance can extend temporal integration and support memory across stimulus-free intervals^16–19^. Yet experimental evidence establishing a causal role for defined inhibitory microcircuits in maintaining persistent activity during working memory behavior is limited. Addressing this question requires a system in which identified circuit elements can be precisely monitored and perturbed during behavior.

*Drosophila* offers a uniquely tractable system for this purpose, allowing cell-type-specific manipulation of defined microcircuits and the simultaneous recording of their activity during learning^20^. Flies are capable of trace conditioning^21–23^, and previous work identified persistent calcium dynamics in GABAergic ellipsoid body (EB) ring neurons during the trace interval, dynamics absent during delay conditioning^24^. These observations suggest that persistent EB activity is selectively engaged when working memory is required. However, whether reciprocal inhibitory interactions within defined EB microcircuits are required to sustain persistent neural representations across the trace interval remains unknown^25^.

Here, we address this question by combining behavior, targeted neural circuit perturbations, and *in vivo* imaging of both calcium and real-time neurotransmitter dynamics. We identify a reciprocal inhibitory loop within the EB that is selectively required to maintain neural representations across the trace interval in trace learning, but dispensable for delay learning, demonstrating that excitation alone cannot support temporal maintenance. We further show that glutamatergic co-transmission and nitric oxide signaling enhance inhibitory efficacy and shape the temporal dynamics of circuit activity. Together, these findings define an inhibitory-modulatory microcircuit that stabilizes persistent neural activity during working memory-dependent behavior, providing a mechanistic framework in which inhibition plays a central role in working memory generation and maintenance.

### Inhibition is necessary for CS-US association in trace conditioning

In the fly central complex, the EB substructure is comprised of a group of neurons that extend their axons to form multiple concentric rings^26^. The aforementioned ER2/4m subpopulation, found to be mobilized during visual conditioning^21^, reciprocate synapses heavily with the ER3/4d neural population, that are also GABAergic in nature^27,28^ (Fig. 1a-c). Here, we questioned whether these two ring neuron subgroups within the central complex formed an inhibitory microcircuit driving neural persistence and working memory observed specifically during trace (but not delay) conditioning (Fig. 1d).

**Figure 1.**
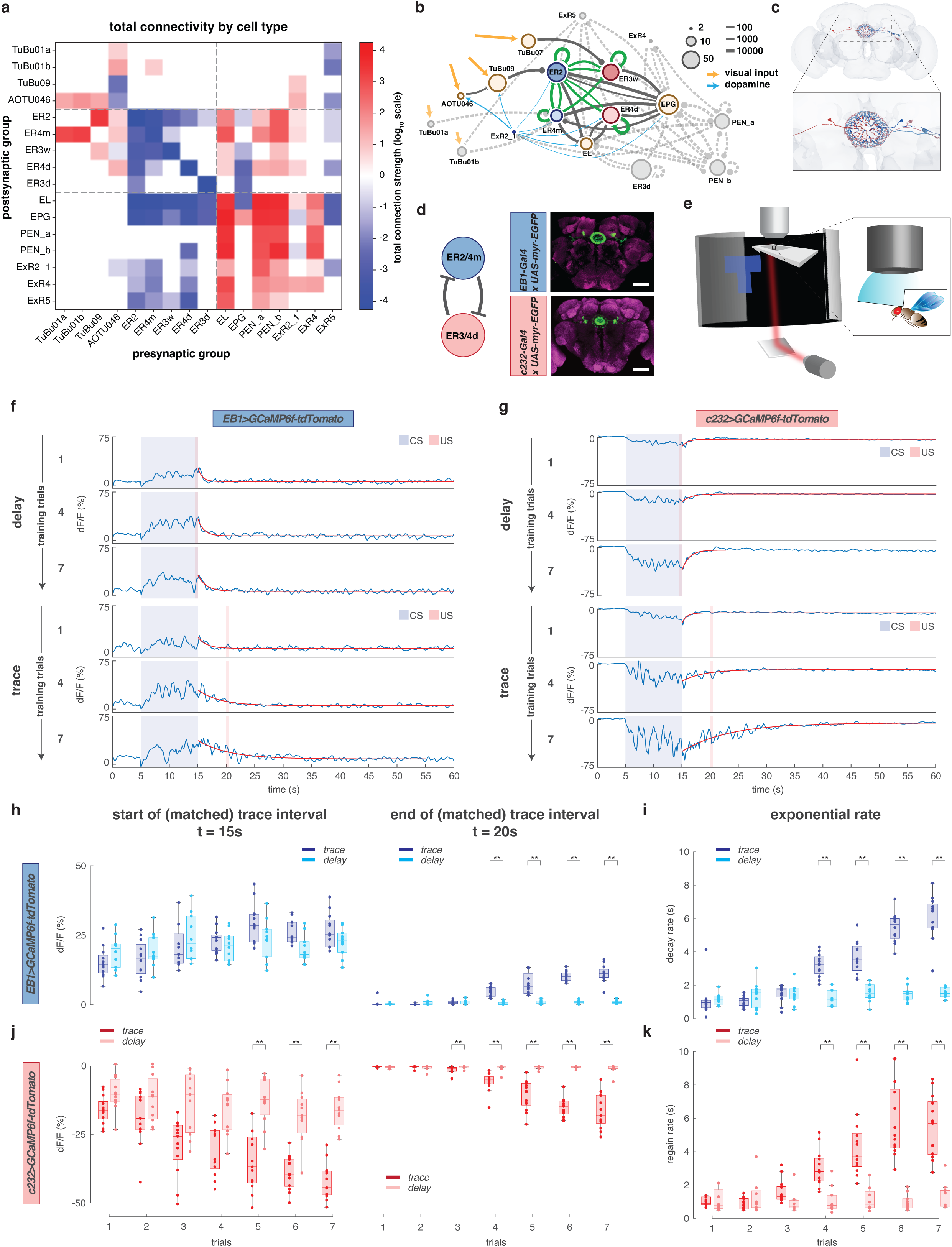
Inhibitory subunits are task-selective and exhibit opposing neuronal dynamics during working memory processing. **(a)** Connectivity matrix by cell type, representing the total numbers of synapses between each pair of neuronal groups. Connection strengths are mapped with a sign function according to neurotransmitter type, and values are shown on a logarithmic scale (see methods, fly connectome analysis). **(b)** Network connectivity graph of dominant excitatory and inhibitory connectivity patterns of the population. Dominant connectivity was established by thresholding the total connectivity strengths on the logarithmic scale; all transformed values below this threshold are not shown to highlight the prominent network structure and major anatomically connected partners of EB ER2/4m neurons. The widths of the edges are drawn in proportion to connectivity strength. Recurrent connections among the EB ring neurons are shown in green. Node sizes were set proportional to the logarithmic scale of the number of neurons within each cell type. **(c)** 3D visualization of 4 select neuron members of ER2, ER4m, ER3w, ER4d in dark blue, light blue, dark red, light red, respectively. This panel is reproduced from FlyWire anatomical reconstruction. Inset: zoomed-in display of synaptic contact sites in the EB forming a ring structure. **(d)** Expression pattern of female EB ER2/4m neurons (*EB1-Gal4>>UAS-myr-EGFP*), and (h) EB ER3/4d neurons (*c232-Gal4>>UAS-myr-EGFP*). Scale bars are 50 μm, n = 3 brains immunostained per genotype. **(e)** Visual conditioning assay coupled with two-photon *in vivo* brain imaging - tethered fly under an objective (inset) shown an upright-T paired with heat. **(f)** Ratiometric imaging of *EB1-Gal4>>UAS-GCaMP6f*.*myr-tdTomato* female during delay conditioning (top) and trace conditioning (bottom). Shown, dF_ratio_/F_ratio_ activity (trials 1, 4, 7). Single-term exponential curve fits (red) through dF_ratio_/F_ratio_ activity starting at CS offset (see methods, fluorescence quantification). **(g)** Ratiometric imaging of *c232-Gal4>>UAS-GCaMP6f*.*myr-tdTomato* female during delay conditioning (top) and trace conditioning (bottom). Shown, dF_ratio_/F_ratio_ activity (trials 1, 4, 7). Single-term exponential curve fits (red) through dF_ratio_/F_ratio_ activity starting at CS offset. **(h)** dF_ratio_/F_ratio_ magnitude (y-intercept) of the exponential fit of ER2/4m (*EB1-Gal4>>UAS-GCaMP6f*.*myr-tdTomato*) calcium activity for delay conditioning (light blue, n = 10 flies) vs. trace conditioning (dark blue, n = 12 flies) at (left) CS offset (start of trace interval, t = 15 s) and (right) 5 s after CS offset (end of trace interval in trace conditioning, t = 20 s). **(i)** ER2/4m decay constants (tau) of exponential fit of dF_ratio_/F_ratio_ calcium activity for delay conditioning (light blue) vs. trace conditioning (dark blue). Tau is defined as the amount of time the activity signal would take to decay/grow by a factor of 1/e (see methods, exponential model fitting). **(j)** dF_ratio_/F_ratio_ magnitude (y-intercept) of the exponential fit of ER3/4d (*c232-Gal4>>UAS-GCaMP6f*.*myr-tdTomato*) calcium activity for delay conditioning (light red, n = 12 flies) vs. trace conditioning (dark red, n = 13 flies) at (left) CS offset (start of trace interval, t = 15 s) and (right) US onset (end of trace interval, t = 20 s). **(k)** ER3/4d growth constants (tau) of exponential fit of dF_ratio_/F_ratio_ calcium activity for delay conditioning (light red) vs. trace conditioning (dark red). Boxplot center (median), edges (IQR), whiskers (1.5x IQR). Scatters represent single-fly metrics. Groups compared using two-factor ART-ANOVA. ** indicates p-value < 0.01.

The conditioning paradigms involved presentation of a visual T-shaped image (upright or inverted) in front of the fly (CS) followed by a heat punishment (US), as has been described previously (Fig. 1e)^21^. Delay conditioning involved presentation of the US in the final moments of the CS, with both ending together. This CS-US coincidence is absent in trace conditioning as it included a 5 s trace interval between the end of the CS and presentation of US. To characterize the trial-by-trial, real-time calcium dynamics of these individual neurons during training, we performed *in vivo* two-photon ratiometric calcium imaging, expressing calcium-dependent GcaMP6f and calcium-independent tdTomato reporters specifically in either EB ER2/4m (*EB1-Gal4>>UAS-GcaMP6f*.*myr-tdTomato*) or ER3/4d *(c232-Gal4>> UAS-GcaMP6f.myr-tdTomato)* ring neurons, during trace and delay conditioning (Fig. 1f, g).

During trace conditioning, ER2/4m neurons exhibited calcium transients that progressively increased in intensity and oscillatory dynamics with repeated training, especially during the trace interval (Fig. 1h, Extended Data Fig. 1). Further, repeated training produced an increase in calcium signal persistence (slower decay following CS termination) in the ER2/4m neurons, a signature that emerged during trace conditioning but was not observed during delay conditioning (Fig. 1i, Extended Data Fig. 2). In contrast, ER3/4d neural activity appeared to run counter to ER2/4m neurons during both delay and trace conditioning (Fig. 1g, Extended Data Fig. 3,4). In initial training trials, calcium activity in ER3/4d neurons decreased during CS presentation in both trace and delay conditioning (Fig. 1j, Extended Data Fig. 3, 4). However, as trace conditioning progressed, this suppression grew stronger during the CS, continuing through the trace interval, a training-dependent effect not observed in delay conditioning (Fig. 1k).

Together, these results indicate an inverse relationship between these two interneuron populations, suggesting that they play opposing complementary “circuit” roles that support the neural synchrony required to maintain the visual trace during the trace interval.

To confirm the necessity of successful trace learning requiring the selective activation of the ER2/4m neural population and suppression of ER3/4d, during the trace interval, we optogenetically activated each of these inhibitory neuron populations using a red-light-activating laser pulse during the (matched) trace interval at 17.5 s (Fig. 2a). Activating ER2/4m neurons (*EB1-Gal4>>UAS-CsChrimson*) resulted in a modest improvement in trace learning (Fig. 2b - left), whereas activating ER3/4d neurons (*c232-Gal4>>UAS-CsChrimson*) led to a drastic decrease in trace learning (Fig. 2b, right). As anticipated, neither manipulation affected delay learning. While ER2/4m and ER3/4d subgroups are selectively active in both trace and delay conditioning, the trace interval elicits a uniquely sustained activity that raises questions about how these inhibitory-to-inhibitory interactions shape the activity underlying trace conditioning.

**Figure 2.**
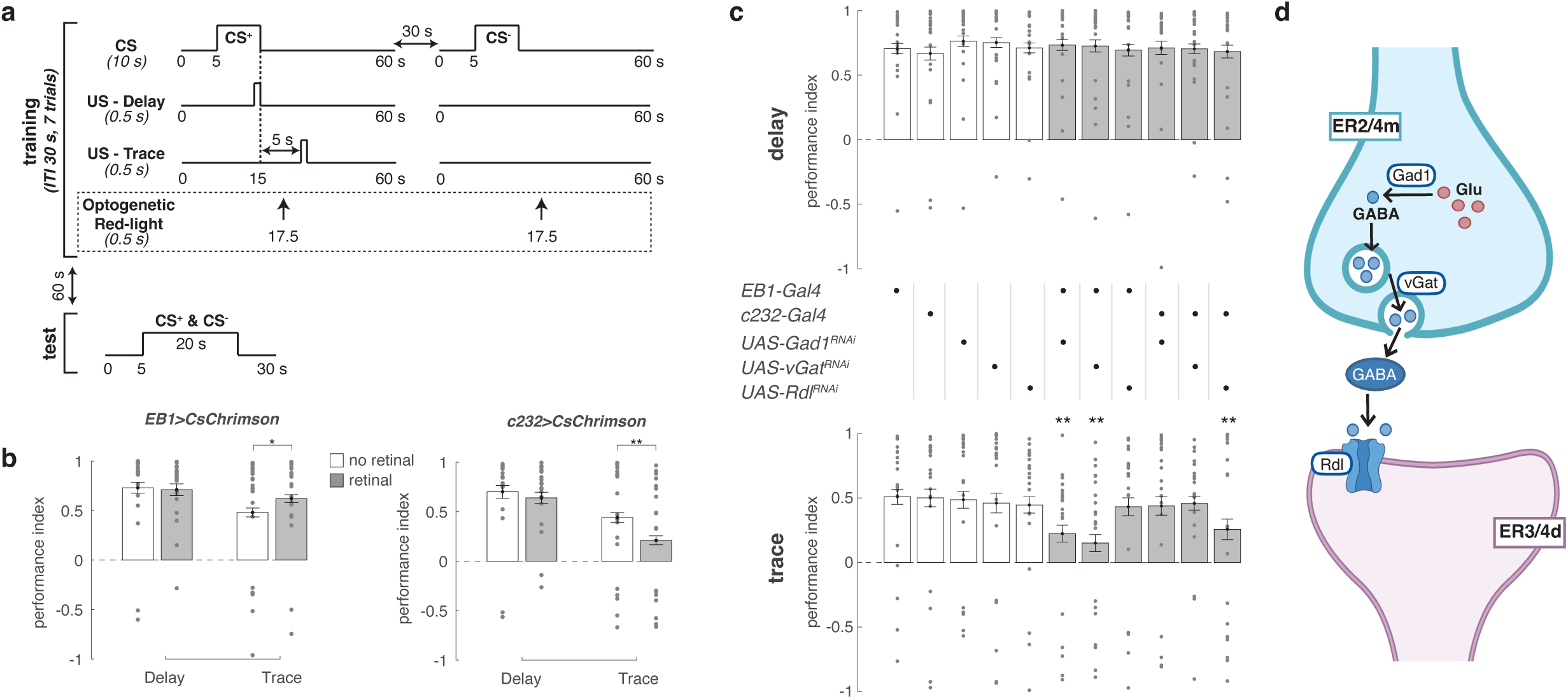
Feedforward GABAergic inhibition of ellipsoid body ER3/4d neurons from ER2/4m neurons is required for trace, and not delay, learning. **(a)** Delay or trace conditioning protocol wherein CS^+^ (either an upright- or inverted-T) was paired with US (heat), and CS^-^ (upright- or inverted-T, whichever not used as CS^+^) without US. Red-light pulse was used for *CsChrimson* driven optogenetic activation of targeted neurons during training. Post-training, fly’s CS^+^ vs. CS^-^ closed-loop orientation was scored in a single 20 s test. **(b)** Delay and trace learning (5 s TI) PI, mean with s.e.m., n = 20 flies per group, on standard food (white) and all-trans-retinal supplemented food (gray), where ER2/4m (*EB1-Gal4>>UAS-CsChrimson*) (left), and ER3/4d (*c232-Gal4>>UAS-CsChrimson*) ellipsoid body ring neurons (right), are optogenetically stimulated as shown in (a). Groups compared using unpaired two-sided Mann–Whitney U tests. **(c)** Delay conditioning (top row) vs. trace conditioning (5 s TI, bottom row) PI (mean with s.e.m., n = 20 flies per group) with impaired Gad1, vGat, Rdl receptor signaling in EB ER2/4m, ER3/4d neurons. Experimental genotypes (gray) compared to parentals using Kruskall-Wallis and post-hoc bonferroni-corrected unpaired two-sided Mann–Whitney U tests. **(d)** Illustration of the GABA signaling system at play during trace conditioning in which GABA is transmitted from ER2/4m neurons to ER3/4d neurons, as evidenced by the RNAi knockdown loss-of-function behavioral results in (c). In (b) and (c), scatters represent single-fly PI. * and ** indicate p-values < 0.05, and 0.01 respectively.

### ER2/4m feedforward GABAergic signaling influences persistent calcium dynamics

To understand the underlying molecular mechanisms of inhibitory interactions between ER2/4m and ER3/4d neurons and their influence on signal strength, persistence, and oscillatory dynamics, we knocked down several classes of genes considered to be necessary for inhibition between these neural populations. We first selectively targeted GABAergic transmission by disrupting its production (*Gad1* GABA-synthesizing protein) and transport (*Vgat* vesicular glutamate transporter), as well as its reception (*Rdl* GABA_A_ receptor) in ER2/4m and ER3/4d neurons^29^. Behavioral data showed that RNAi-based knockdowns of both GABA production (*UAS-Gad1^RNAi^*) and transport (*UAS-Vgat^RNAi^*) in ER2/4m neurons impaired behavioral performance in trace, but not delay, learning (Fig. 2c). Additionally, RNAi knockdown of GABA_A_ receptor (*UAS-Rdl^RNAi^*) in ER3/4d neurons also compromised trace, but not delay, learning (Fig. 2c). Strikingly, disrupting reciprocal GABA transmission from ER3/4d neurons and GABA_A_ reception in ER2/4m neurons did not affect trace or delay learning. An intriguing result, since activating the ER3/4d population impaired trace learning (Fig. 2b - right), suggesting that GABAergic inhibition from ER3/4d neurons may be dispensable, or that these neurons influence learning through another form of inhibitory communication. Together, these results do implicate feedforward GABAergic transmission from ER2/4m to ER3/4d neurons as necessary for mediating inhibitory-to-inhibitory neural communication during trace learning (Fig. 2d).

Based on these behavioral data, we hypothesized that sustained inhibition from ER2/4m neurons is necessary for maintaining signaling stability during the trace interval and driving neural synchrony. We tested this rationale in two ways. First, by performing ratiometric calcium imaging of ER2/4m neurons with an RNAi knockdown of GABA synthesis in those same neurons (*EB1-Gal4>>UAS-GcaMP6f*.*myr-tdTomato, UAS-Gad1^RNAi^*) and second, ratiometric calcium imaging of ER3/4d neurons with an RNAi knockdown of GABA reception (*c232-Gal4>> UAS-GcaMP6f.myr-tdTomato, UAS-Rdl^RNAi^*), respectively (Fig. 3a, d). During trace conditioning, RNAi knockdown of GABA synthesis in ER2/4m neurons largely spared activity during CS (Fig. 3b) but disrupted the characteristic persistence (Fig. 3b, c) and frequency increase (Extended Data Fig. 1d) normally observed during the trace interval. These effects were specific to trace conditioning and expectedly, were not observed during delay conditioning (Extended Data Fig. 2). Similarly, when reducing postsynaptic GABAAR availability in ER3/4d neurons, the observed suppression during CS and trace interval was lost (Fig. 3e), resulting in a loss of signal decay and consequently regain (Fig. 3e, f, Extended Data Fig. 3). Likewise, during delay conditioning, this knockdown also reduced CS-based suppression (Extended Data Fig. 4), yet behavior remained unaffected, emphasizing that inhibitory signaling, though observed, is not required for delay conditioning^24^. These findings suggest that both GABA synthesis in ER2/4m neurons as well as GABA reception in ER3/4d neurons play a critical role in maintaining the CS neural representation during the trace interval and supporting trace conditioning, yet we still lack insight into how reciprocal communication from ER3/4d is mediated.

**Figure 3.**
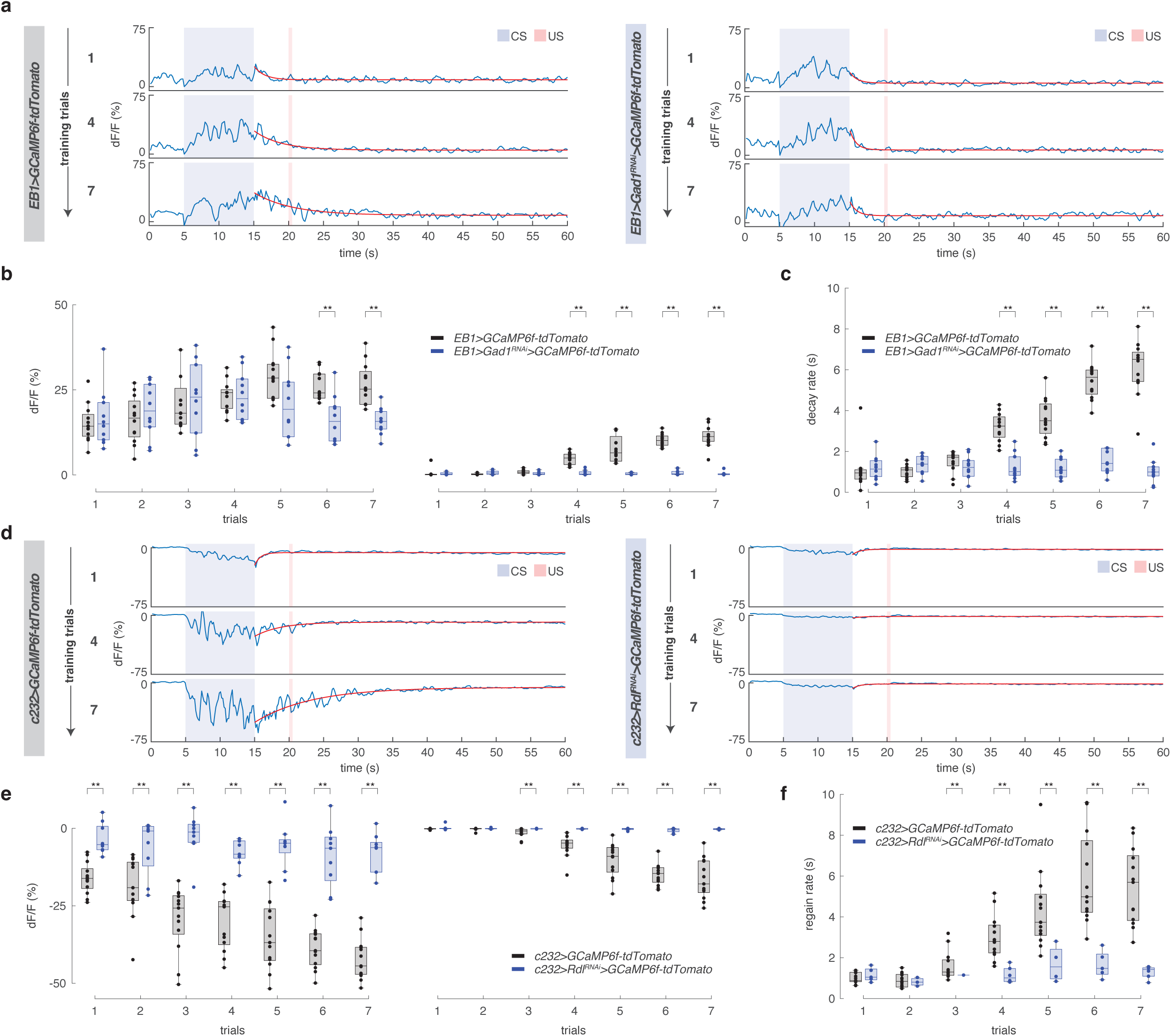
Sustained neural stability requires coordinated, feedforward GABAergic inhibition of ellipsoid body ER3/4d from ER2/4m neurons. **(a)** Ratiometric imaging of *EB1-Gal4>>UAS-GCaMP6f*.*myr-tdTomato* (left) and *EB1-Gal4>>UAS-GCaMP6f*.*myr-tdTomato,UAS-Gad1^RNAi^* (right) females during trace conditioning. Shown, dF_ratio_/F_ratio_ activity (trials 1, 4, 7). Single-term exponential curve fits (red) through dF_ratio_/F_ratio_ activity starting at CS offset (see methods, fluorescence quantification). **(b)** dF_ratio_/F_ratio_ magnitude (y-intercept) of the exponential fit of ER2/4m calcium activity for *EB1-Gal4>>UAS-GCaMP6f*.*myr-tdTomato* (black, n = 12 flies) and *EB1-Gal4>>UAS-GCaMP6f*.*myr-tdTomato,UAS-Gad1^RNAi^* (blue, n = 10 flies) at (left) CS offset (start of trace interval, t = 15 s) and (right) US onset (end of trace interval, t = 20 s). **(c)** ER2/4m decay constants (tau) of exponential fit of dF_ratio_/F_ratio_ calcium activity for *EB1-Gal4>>UAS-GCaMP6f*.*myr-tdTomato* (black) and *EB1-Gal4>>UAS-GCaMP6f*.*myr-tdTomato,UAS-Gad1^RNAi^* (blue). Tau is defined as the amount of time the activity signal would take to decay/grow by a factor of 1/e (see methods, exponential model fitting). **(d)** Ratiometric imaging of *c232-Gal4>>UAS-GCaMP6f*.*myr-tdTomato* (left) and *c232-Gal4>>UAS-GCaMP6f*.*myr-tdTomato,UAS-Rdl^RNAi^*(right) females during trace conditioning. Shown, dF_ratio_/F_ratio_ activity (trials 1, 4, 7). Single-term exponential curve fits (red) through dF_ratio_/F_ratio_ activity starting at CS offset. **(e)** dF_ratio_/F_ratio_ magnitude (y-intercept) of the exponential fit of ER3/4d calcium activity for *c232-Gal4>>UAS-GCaMP6f*.*myr-tdTomato* (black, n = 13 flies) and *c232-Gal4>>UAS-GCaMP6f*.*myr-tdTomato,UAS-Rdl^RNAi^* (blue, n = 9 flies) at (left) CS offset (start of trace interval, t = 15 s) and (right) US onset (end of trace interval, t = 20 s). **(f)** ER3/4d growth constants (tau) of exponential fit of dF_ratio_/F_ratio_ calcium activity for *c232-Gal4>>UAS-GCaMP6f*.*myr-tdTomato* (black) and *c232-Gal4>>UAS-GCaMP6f*.*myr-tdTomato,UAS-Rdl^RNAi^* (blue). Boxplot center (median), edges (IQR), whiskers (1.5x IQR). Scatters represent single-fly metrics. Groups compared using two-factor ART-ANOVA. ** indicates p-value < 0.01.

### Nitric oxide and glutamate support persistent activity in a reciprocal inhibitory circuit

Reciprocal inhibitory signaling is being increasingly recognized as key to supporting the brain’s adaptability by modulating the temporal structure and synchrony of neural activity^30,31^. To explore the possibility of crosstalk from ER3/4d to ER2/4m neurons beyond GABAergic transmission, we investigated the potential role of nitric oxide (NO) given its involvement in *Drosophila* spatial memory^32–34^. NO can serve as a gasotransmitter, activating sGC and cGMP biosynthesis^35^, that can ultimately alter cellular function of target neurons locally near the source of its production^36,37^. Here, we sought to determine whether the production and release of NO might result in recurrent communication to influence trace learning. Using an RNAi knockdown approach, we blocked nitric-oxide synthase (*UAS-dNOS^RNAi^*) signaling and tested for effects on trace and delay learning. Notably, we found blocking NO production from ER3/4d neurons selectively impaired trace, not delay, conditioning, while disrupting NO synthase in ER2/4m neurons did not affect either learning paradigm (Fig. 4a). This led us to question what might trigger NO synthase in ER3/4d neurons. A potential insight may be derived from the possibility that *Drosophila* GABAergic neurons, similar to mammals^38,39^, can use glutamate as a co-transmitter^40^, the reception of which could trigger nitric oxide (NO) release in neighboring neurons^41^. We therefore tested for this possibility, by assaying for behavioral changes due to blocking vesicular glutamate transport^42^ (*UAS-dVglut^RNAi^*) using the RNAi knockdown approach. Behavioral results revealed that disrupting *dVglut* signaling in ER2/4m neurons selectively impaired trace learning, leaving delay learning intact (Fig. 4a). Further, this effect was specific to blocking glutamate transport from ER2/4m neurons, as disrupting *dVglut* signaling in ER3/4d neurons had no deleterious effect on trace or delay learning (Fig. 4b).

**Figure 4.**
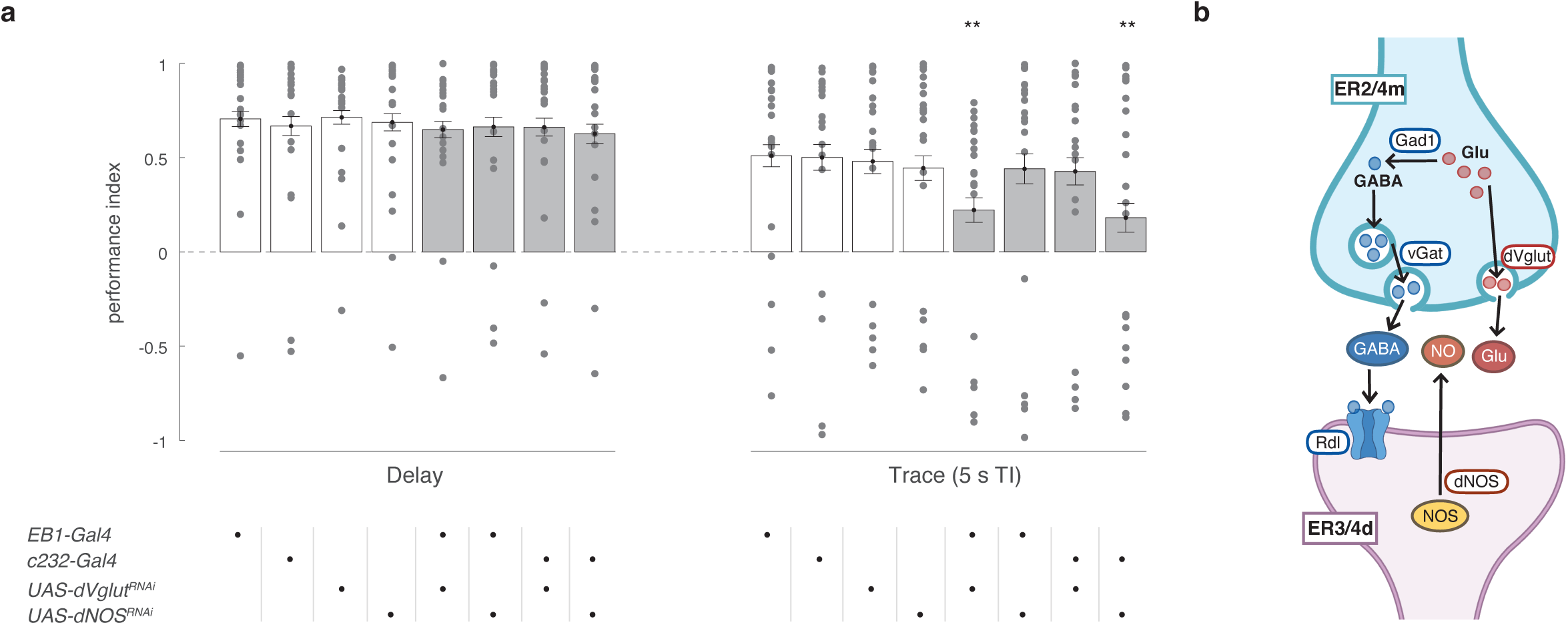
Co-transmission of glutamate from ER2/4m neurons and nitric oxide synthase-mediated signaling from ER3/4m neurons is required for trace, and not delay, learning. **(a)** Delay conditioning (left) and trace conditioning (5 s TI, right) PI (mean with s.e.m., n = 20 flies per group) with impaired dVglut, dNOS receptor signaling in EB ER2/4m, ER3/4d neurons. Experimental genotypes (gray) compared to parentals using Kruskall-Wallis and post-hoc bonferroni-corrected unpaired two-sided Mann–Whitney U tests. Scatters represent single-fly PI. ** indicates p-value < 0.01. **(b)** Illustration of Glutamate/GABA co-signaling system at play during trace conditioning in which Glu and GABA are co-transmitted from ER2/4m neurons, and NO is produced by ER3/4d neurons, as evidenced by the RNAi knockdown loss-of-function behavioral results in (a).

Since behavioral impairment implicated a Glu/NO signaling pathway underlying trace learning, we performed ratiometric calcium imaging of ER3/4d neurons with an RNAi knockdown of NO synthesis in those same neurons (*c232-Gal4>>UAS-GcaMP6f*.*myr-tdTomato, UAS-dNOS^RNAi^*), to examine the effects on calcium signaling dynamics from this secondary signaling pathway (Fig. 5a). During trace conditioning, knockdown of NO synthesis in ER3/4d neurons resulted in a weakened signal decrease compared to wildtype, but stronger than GABA_A_R knockdown activities (Fig. 5b, Extended Data Fig. 3). This weakened response suggests that the underlying mechanism driving ER3/4d activity suppression remains partly functional without NO during CS, however, appears insufficient to drive sustained suppression through the trace interval, rendering it incapable of trace learning (Fig. 5c). Calcium dynamics during delay conditioning remained intact despite this knockdown (Extended Data Fig. 4), highlighting that this secondary signaling mechanism is dispensable for delay conditioning, in contrast to the more cognitively demanding trace conditioning.

**Figure 5.**
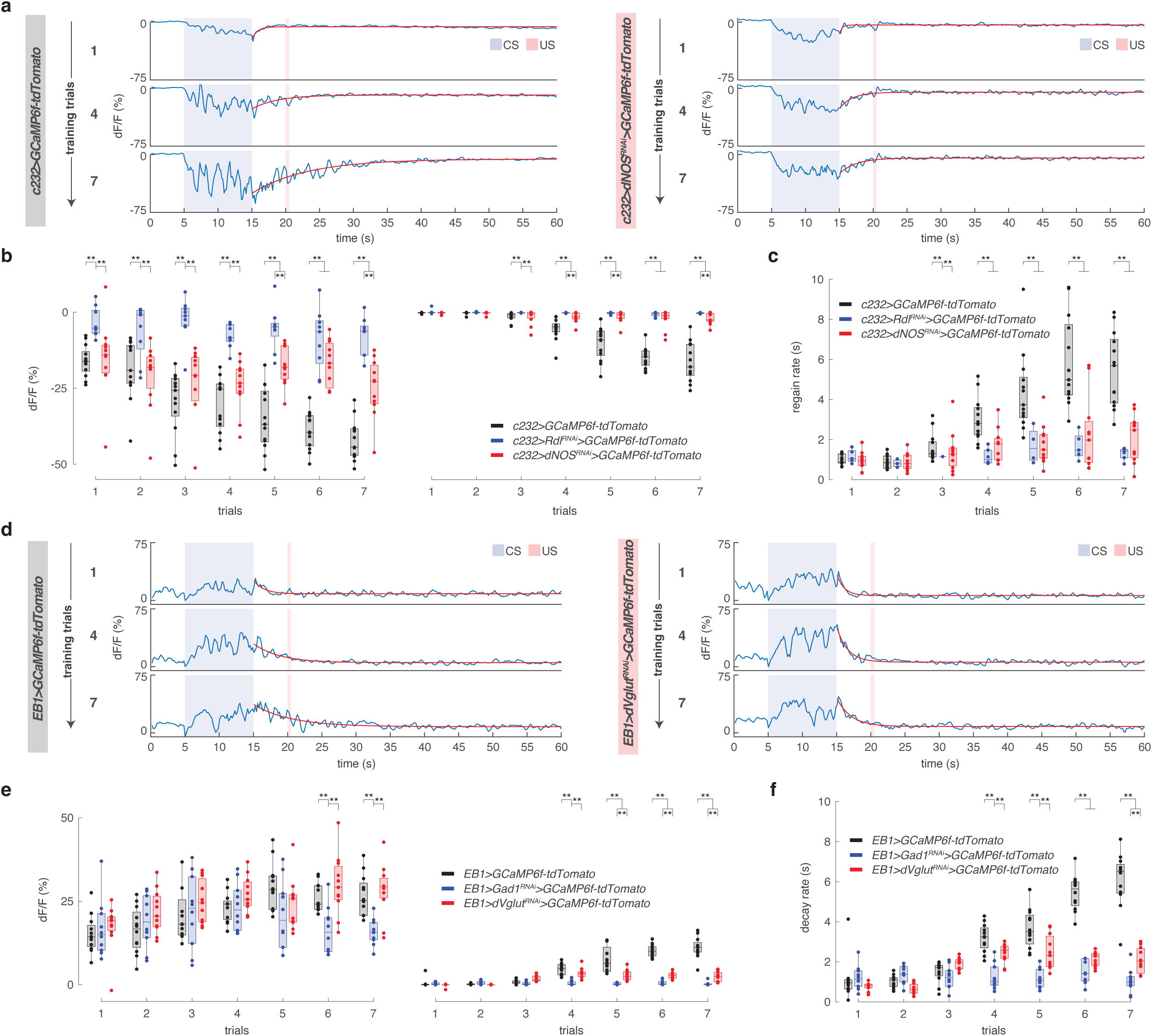
Supportive role of glutamate and nitric oxide in strengthening neural persistence during trace conditioning. **(a)** Ratiometric imaging of *c232-Gal4>>UAS-GCaMP6f*.*myr-tdTomato* (left) and *c232-Gal4>>UAS-GCaMP6f*.*myr-tdTomato,UAS-dNOS^RNAi^*(right) females during trace conditioning. Shown, dF_ratio_/F_ratio_ activity (trials 1, 4, 7). Single-term exponential curve fits (red) through dF_ratio_/F_ratio_ activity starting at CS offset (see methods, fluorescence quantification). **(b)** dF_ratio_/F_ratio_ magnitude (y-intercept) of the exponential fit of ER3/4d calcium activity for *c232-Gal4>>UAS-GCaMP6f*.*myr-tdTomato* (black, n = 13 flies), *c232-Gal4>>UAS-GCaMP6f*.*myr-tdTomato,UAS-Rdl^RNAi^*(blue, n = 9 flies), and *c232-Gal4>>UAS-GCaMP6f*.*myr-tdTomato,UAS-dNOS^RNAi^* (red, n = 11 flies) at (left) CS offset (start of trace interval, t = 15 s) and (right) US onset (end of trace interval, t = 20 s). **(c)** ER3/4d growth constants (tau) of exponential fit of dF_ratio_/F_ratio_ calcium activity for *c232-Gal4>>UAS-GCaMP6f*.*myr-tdTomato* (black), *c232-Gal4>>UAS-GCaMP6f*.*myr-tdTomato,UAS-Rdl^RNAi^*(blue), and *c232-Gal4>>UAS-GCaMP6f*.*myr-tdTomato,UAS-dNOS^RNAi^*(red). Tau is defined as the amount of time the activity signal would take to decay/grow by a factor of 1/e (see methods, exponential model fitting). **(d)** Ratiometric imaging of *EB1-Gal4>>UAS-GCaMP6f*.*myr-tdTomato* (left) and *EB1-Gal4>>UAS-GCaMP6f*.*myr-tdTomato,UAS-dVglut^RNAi^*(right) females during trace conditioning. Shown, dF_ratio_/F_ratio_ activity (trials 1, 4, 7). Single-term exponential curve fits (red) through dF_ratio_/F_ratio_ activity starting at CS offset. **(e)** dF_ratio_/F_ratio_ magnitude (y-intercept) of the exponential fit of ER2/4m calcium activity for *EB1-Gal4>>UAS-GCaMP6f*.*myr-tdTomato* (black, n = 12 flies), *EB1-Gal4>>UAS-GCaMP6f*.*myr-tdTomato,UAS-Gad1^RNAi^*(blue, n = 10 flies), and *EB1-Gal4>>UAS-GCaMP6f*.*myr-tdTomato,UAS-dVglut^RNAi^* (red, n = 11 flies) at (left) CS offset (start of trace interval, t = 15 s) and (right) US onset (end of trace interval, t = 20 s). **(f)** ER2/4m decay constants (tau) of exponential fit of dF_ratio_/F_ratio_ calcium activity for *EB1-Gal4>>UAS-GCaMP6f*.*myr-tdTomato* (black), *EB1-Gal4>>UAS-GCaMP6f*.*myr-tdTomato,UAS-Gad1^RNAi^*(blue), and *EB1-Gal4>>UAS-GCaMP6f*.*myr-tdTomato,UAS-dVglut^RNAi^*(red). Boxplot center (median), edges (IQR), whiskers (1.5x IQR). Scatters represent single-fly metrics. Groups compared using two-factor ART-ANOVA. ** indicates p-value < 0.01.

To address how possible glutamate transmission from ER2/4m neurons influence neural activity dynamics, we monitored calcium transients in flies with RNAi knockdown of *dVglut* signaling in ER2/4m neurons (*EB1-Gal4>>UAS-GcaMP6f.myr-tdTomato, UAS-dVglut^RNAi^*) (Fig. 5d). During trace conditioning, *dVglut*-impaired flies exhibited an increase in calcium activity, frequency, and persistence (slowing decay) during the initial training trials, however, that increase plateaued below wildtype levels as trials progressed (Fig. 5fe, Extended Data Fig. 1d).

During delay conditioning, *dVglut*-impaired flies showed calcium dynamics similar to wildtype, indicating that delay conditioning-related calcium dynamics are preserved despite the impaired glutamate transport (Extended Data Fig. 2). This result specifically points to the importance of glutamate release in maintaining the CS neural representation in ER2/4m neurons during the trace interval, uncovering an inhibitory reciprocal network underlying the observed sustained activity. Taken together, these results suggest that ER2/4m and ER3/4d neurons maintain neural synchrony through a reciprocal signaling cascade in which glutamate and GABA are co-transmitted differentially by ER2/4m neurons and NO is produced by ER3/4d neurons.

### Glutamatergic neuromodulation increases GABAergic inhibitory efficacy and signal persistence

Co-transmission of glutamate and GABA has been widely investigated in mammals^43^ and insects^44^, but how these signaling dynamics influence working memory have yet to be identified. Here, we sought to directly monitor glutamate and GABA signaling from ER2/4m neurons using real-time fluorescent indicators iGluSnFR (*EB1-Gal4>>UAS-iGluSnFR*) and iGABASnFR2 (*EB1-Gal4>>UAS-iGABASnFR2*) respectively, and to examine whether these neurochemicals displayed distinct spatiotemporal activity patterns during delay and trace conditioning.

During both forms of conditioning, we observed a sharp rise in glutamate activity in ER2/4m neurons after US presentation (Fig. 6a, Extended Data Fig. 5a). This response progressively diminished in both paradigms (Fig. 6b, Extended Data Fig. 5b), however, specifically during trace conditioning, we observed the phasic glutamate activity shift from after the US towards the trace interval (Fig. 6c, Extended Data Fig. 5c). Further, we observed three notable quantitative neural signatures of the spatiotemporal patterns of glutamate activity during trace conditioning. First, glutamate levels during the trace interval appeared to resemble an inverted U-shaped response, non-existent initially, emerging around trial 3, peaking around trial 5, and subsequently declining (Fig. 6b, e, Extended Data Fig. 5b), a phenomenon studied extensively in both humans and higher order mammals^45,46^. Second, in initial training trials, the observed glutamate activity post-US did not coincide with increased calcium signaling, indicating engagement of a calcium-independent signaling cascade (Fig. 6d). Third, the rise in glutamate activity precedes the emergence of prolonged ER2/4m calcium activity that is observed a trial later (Fig. 6e). Finally, during delay conditioning, a weaker glutamate response is evoked post-US than what was observed in trace conditioning (Extended Data Fig. 5).

**Figure 6.**
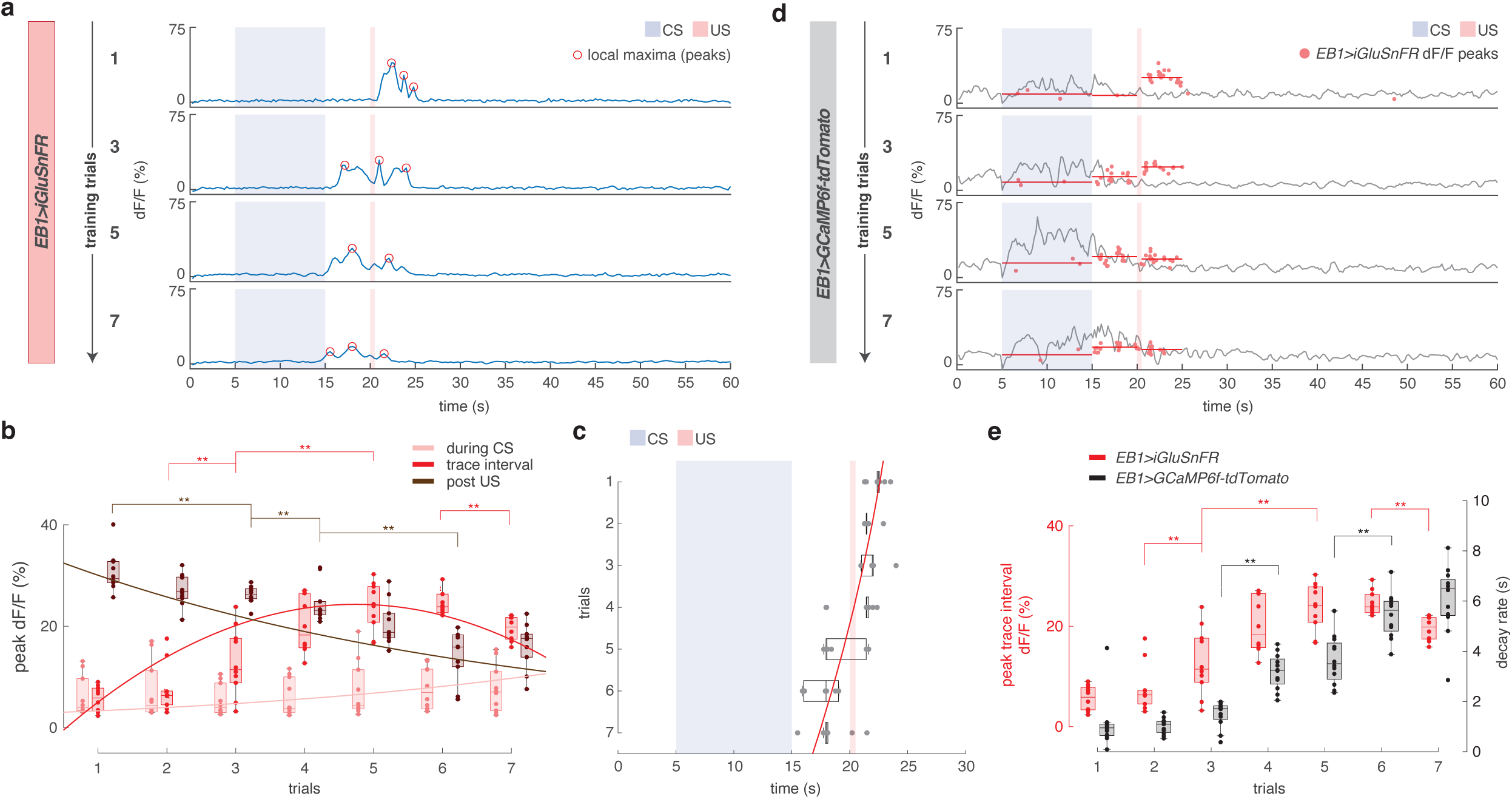
Phasic glutamatergic release shifts towards the trace interval precede strengthened neural persistence during trace conditioning. **(a)** *In vivo* fluorescent glutamate indicator imaging from ER2/4m neurons, *EB1-Gal4>>UAS-iGluSnFR* female during trace conditioning. Shown, dF/F activity (trials 1, 3, 5, 7) with local maxima peaks (red circles) of dF/F activity (see methods, fluorescence quantification, and GluSnFR peak detection analysis). **(b)** Peak dF/F activity during CS (light red), during TI (red), and post-US (dark red) of trace conditioning (n = 10 flies). **(c)** Timing of the highest local maxima peak of dF/F activity over trials during trace conditioning. In (b) and (c), quadratic (second degree) polynomial curve fits through median dF/F activity. **(d)** Scatters of local maxima peaks of dF/F activity for *EB1-Gal4>>UAS-iGluSnFR* (red, n = 10 flies) with median activity levels (during CS, during TI, and post US), overlaid on a ratiometric calcium imaging of *EB1-Gal4>>UAS-GCaMP6f*.*myr-tdTomato* female during trace conditioning. Shown, dF_ratio_/F_ratio_ activity (trials 1, 3, 5, 7). **(e)** Peak dF/F ER2/4m glutamate release activity during TI of trace conditioning for *EB1-Gal4>>UAS-iGluSnFR* (red, n = 10 flies), from (b), against ER2/4m decay constant (tau) of exponential fit of dF_ratio_/F_ratio_ calcium activity for *EB1-Gal4>>UAS-GCaMP6f*.*myr-tdTomato* (black, n = 12 flies). Tau is defined as the amount of time the activity signal would take to decay/grow by a factor of 1/e (see methods, exponential model fitting). Boxplot center (median), edges (IQR), whiskers (1.5x IQR). Scatters represent single-fly metrics. Groups compared using two-factor ART-ANOVA. ** indicates p-value < 0.01.

In contrast to glutamate, GABA increases in ER2/4m neurons first emerge during CS presentation and rapidly decline after CS offset in both trace (Fig. 7a, Extended Data Fig. 6) and delay conditioning (Extended Data Fig. 7). However, as trials progress, GABA activity becomes more sustained through the trace interval, a pattern not observed during delay conditioning.

**Figure 7.**
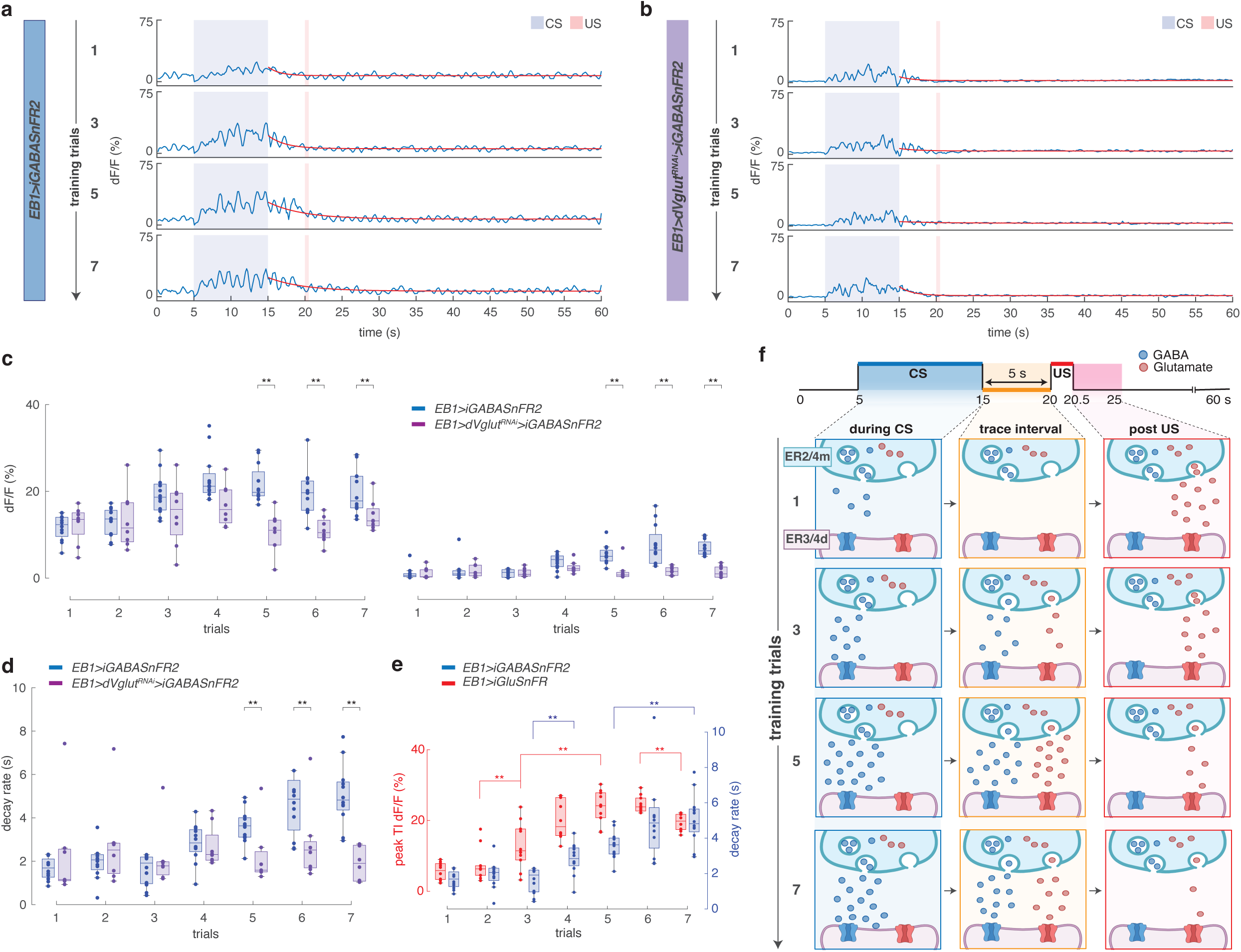
Precise temporal alignment between GABAergic signaling and glutamatergic release from ER2/4m neurons orchestrates underlying observed calcium transients. **(a)** *In vivo* fluorescent GABA indicator imaging from ER2/4m neurons, *EB1-Gal4>>UAS-iGluSnFR2*, and (b) *EB1-Gal4>>UAS-iGABASnFR2,UAS-dVglut^RNAi^*females, during trace conditioning. Shown, dF/F activity (trials 1, 3, 5, 7). Single-term exponential curve fits (red) through dF/F activity starting at CS offset. **(c)** dF/F magnitude (y-intercept) of the exponential fit of ER2/4m GABAergic activity for *EB1-Gal4>>UAS-iGABASnFR2* (blue, n = 12 flies), and *EB1-Gal4>>UAS-iGABASnFR2,UAS-dVglut^RNAi^* (purple, n = 8 flies), at (left) CS offset (start of trace interval, t = 15 s) and (right) US onset (end of trace interval, t = 20 s). **(d)** ER2/4m decay constants (tau) of exponential fit of dF/F GABAergic activity for *EB1-Gal4>>UAS-iGABASnFR2* (blue), and *EB1-Gal4>>UAS-iGABASnFR2,UAS-dVglut^RNAi^* (purple). **(e)** Peak dF/F ER2/4m glutamate release activity during TI of trace conditioning for *EB1-Gal4>>UAS-iGluSnFR* (red, n = 10 flies), from (Fig. 6b, e), against ER2/4m decay constant (tau) of exponential fit of dF/F GABAergic activity for *EB1-Gal4>>UAS-iGABASnFR2* (blue, n = 12 flies), from (d). **(f)** Schematic representation of GABAergic (blue) and glutamatergic (red) neural signaling during the conditioned stimulus (CS), trace interval, and post-unconditioned stimulus (US) period across training trials 1, 3, 5, and 7. Boxplot center (median), edges (IQR), whiskers (1.5x IQR). Scatters represent single-fly metrics. Groups compared using two-factor ART-ANOVA. ** indicates p-value < 0.01.

Because GABA signaling closely mirrored ER2/4m calcium dynamics (Extended Data Fig. 6d), we asked whether glutamate release from ER2/4m was required to reinforce GABAergic activity and initiate the observed signal propagation across the trace interval. *In vivo* imaging of GABA intensity in ER2/4m neurons with glutamate transport knocked down (*EB1-Gal4>>UAS-iGABASnFR2, UAS-dVglut^RNAi^*) revealed reduced activity during CS in later trials (Fig. 7b, c - left). Similarly, blocking glutamate release impaired the characteristic increase and signal persistence seen in GABA activity during the trace interval (Fig. 7c, d), largely mirroring the effect seen in calcium dynamics with a similar knockdown of glutamate release (Fig. 5d-f). Also, much like the calcium signal propagation (Fig. 6e), the rise in glutamate activity preceded the emergence of prolonged ER2/4m GABA activity by a trial (Fig. 7e), suggesting the shift in glutamate activity towards the trace interval is a key factor in initiating and sustaining inhibitory neural persistence. Ultimately, we illustrate that coordination of the time-locked co-release of glutamate and GABA (Fig. 7f) is critical for working memory maintenance in trace conditioning.

## Discussion

Persistent neural activity has long been attributed to recurrent excitation within local circuits^47,48^. Our results instead identify an inhibitory architecture within the EB as the primary driver of persistence during working memory-dependent trace conditioning. Disrupting inhibitory signaling, through mistimed neural activation or selective interference with neurotransmitter signaling, abolished sustained activity and selectively impaired trace, but not delay, learning. These findings demonstrate that excitation alone is insufficient to maintain neural representations across stimulus-free intervals, establishing a causal role for structured inhibitory interactions in generating behaviorally relevant neural persistence *in vivo*.

Crucially, the persistence we observe does not emerge from classical recurrent excitation stabilized by inhibition^49^. It arises from temporally structured, reciprocal inhibitory interactions reinforced by glutamatergic co-transmission and retrograde NO signaling. The source, timing and coordination of transmitter release are central determinants of whether task-relevant activity is sustained. In this circuit, inhibition is not merely restraining excitation, it orchestrates the dynamics that allow activity to bridge temporal gaps.

Mechanistically, ER2/4m neurons co-release GABA and glutamate, and glutamate dynamics were not tightly coupled to ER2/4m calcium influx, consistent with studies highlighting calcium-independent^50^, neuromodulator-triggered release^51^. Extracellular glutamate measurements, as reported by iGluSnFR^52^, may reflect contributions from EB-projecting PPM3 dopaminergic (ExR2) inputs^52,53^ or glial redistribution^54^, however, selectively blocking glutamate release from ER2/4m neurons disrupted both GABA signaling and calcium dynamics, confirming these neurons as a functional source for persistence. Glutamate thus reinforces inhibitory efficacy rather than acting as an independent driver of it^55^.

Retrograde NO signaling from ER3/4d neurons further strengthens this circuit. NO may accumulate across trials to modulate rapid GABA release^56^ and GABA_A_ receptor–associated pathways^57^, enhancing inhibitory gain over behaviorally relevant timescales. Glutamate-mediated facilitation and NO-dependent feedback together close the communication loop, stabilizing activity during trace conditioning. This inhibitory-modulatory motif provides a mechanistic substrate for actively maintained neural representations.

These results have broad implications for working memory models. Persistent activity in the mammalian cortex is often modeled as arising from recurrent excitation, with inhibition balancing network gain^49^. In canonical excitation-inhibition balance networks, recurrent excitation sustains activity while inhibition functions as a stabilizing counterweight that prevents runaway dynamics and preserves persistent firing only when excitation is finely tuned^3^.

However, theoretical work predicts that inhibitory connectivity alone can define network timescales and support transient activity states^16,18,19,58^. Our data provides direct experimental support for this alternative view, reciprocal inhibition can be generative, not simply stabilizing. The selective requirement for inhibition during trace, but not delay, conditioning, argues against models in which inhibition merely scales excitatory persistence, supporting a framework in which inhibitory motifs define the temporal dynamics necessary to bridge stimulus-free intervals. While excitation may still initiate or bias network states, it is the inhibitory architecture that actively maintains them. Thus, working memory maintenance may emerge not from finely tuned excitation, but from inhibitory circuit structure itself.

Effective working memory likely depends on inhibition operating across multiple timescales. Fast, phasic GABAergic signaling provides rapid gating and moment-to-moment control^59,60^, while slower glutamate- and NO-dependent modulation biases network dynamics toward sustained, task-relevant states^61^. Persistence emerges from the interplay between rapid inhibitory gating^60,62^ and slower modulatory reinforcement^62^; without this coordinated interaction^63^, working memory would become either too unstable to retain information or too rigid to update it when necessary.

Together, our findings establish a defined inhibitory microcircuit in which coordinated GABAergic transmission, glutamatergic co-release, and nitric oxide-dependent feedback cooperate to stabilize persistent neural activity during working memory-dependent learning. In this framework, ER2/4m neurons provide sustained inhibitory drive that maintains a conditioned stimulus representation across the trace interval, while glutamate and nitric oxide signaling progressively enhance inhibitory efficacy to reinforce persistence over repeated trials. The selective engagement of this circuit during trace, but not delay, conditioning demonstrates that its activity is contingent upon the requirement to bridge a temporal gap, linking circuit dynamics directly to behavioral demand. By demonstrating that reciprocal inhibitory interactions are necessary for sustaining persistent activity *in vivo*, our results expand the repertoire of circuit architectures capable of supporting working memory and highlight inhibition as an active generator, not merely a stabilizer, of memory traces across time.

## Methods

### Fly stocks

Drosophila melanogaster flies were raised and maintained on a standard cornmeal-agar medium at 25°C, 60% humidity, on a 12-h light –12-h dark cycle. All experimental flies were non-mated females 3-10 days old, isolated post-eclosion at 25°C, and tested 0-3 h before the onset of the dark cycle. Canton-S flies served as the wildtype *Drosophila* strain in this study. Stocks with initials BDSC (and corresponding stock numbers) were obtained from the Bloomington Drosophila Stock Center.

Gal4 drivers for expression in ellipsoid body ring neurons were: ER2/4m, *EB1-Gal4* (BDSC 44409) and ER3/4d, *c232-Gal4* (BDSC 30828). Immunostaining protocol used *UAS-myr-EGFP* (BDSC 32197) in targeted neurons.

Optogenetic activation experiments were carried out using *UAS-CsChrimson*^64^ (BDSC 55136). Newly emerged female flies expressing CsChrimson in targeted Gal4 neurons were collected and raised under controlled conditions in constant darkness at 25°C (60% humidity) on a standard cornmeal medium enriched with 200 μM all-trans-retinal (Sigma-Aldrich). Control sibling flies were also maintained in darkness but received only the standard cornmeal medium without the retinal supplement.

The RNA interference (RNAi) lines used were *UAS-Gad1^RNAi^* (BDSC 28079), *UAS-vGAT^RNAi^*(BDSC 41958), *UAS-Rdl^RNAi^* (BDSC 31286), *UAS-dNOS^RNAi^*(BDSC 28792), and *UAS-dVglut^RNAi^* (BDSC 40845).

We used *20XUAS-GCaMP6f.myr-tdTomato*^65^ (in-house stock as described previously^51^) to perform *in vivo* ratiometric calcium imaging experiments. To monitor activity of glutamate and GABA at pre-synaptic terminals, we used *UAS-iGluSnFR.A184A*^52^ (BDSC 59609) and *UAS-iGABASnFR2*^66^ (gift from G. Turner), respectively.

### Virtual reality tethered-flight behavior assay

Female flies were cold-anesthetized (4°C) and tethered to a stainless steel minutien pin (0.2 mm rod diameter, Fine Science Tools) using UV-cured glue, curing for at least 30 seconds. After a recovery period of at least one hour, these tethered flies were suspended within a high-speed projector-based spherical virtual-reality flight arena. The exterior of the 2 in diameter sphere was coated using a rear-projection medium (Screen Goo, Goo Systems Global) allowing the presentation of dynamic computer-generated stimuli and create an immersive virtual environment.

The tethered-flight arena followed the basic protocol outlined previously^66^ to generate an immersive visual surround. Briefly, a high-speed projector (Texas Instruments, LightCrafter 4500 EVM) projected three virtual views of a scene onto the sphere, the first head on, the other two lateral views reflected off two 2-inch mirrors (Edmund Optics) that were placed on either side to the sphere. This produced a virtual-reality display encompassing around 330° of azimuth and around 85° of elevation. Custom software written in Microsoft Visual C++ using a 3D graphics programming library OpenSceneGraph allowed us to warp and display images on a curved spherical projection screen. The projector was set to display 6-bit images at a resolution of 912 x 1140 at 300 Hz and emitted a blue light which was filtered through a 450-nm long-pass emission filter (62-982, Edmund Optics). Calibration of light intensity was performed using a SpectraScan PR-701S spectroradiometer and the intensity was controlled by altering the current of the projector LED, ultimately set to 10% of the maximum power. The overall radiance inside the spherical area for an all-ON stimulus was set to be 0.4 W m^-2^ sr^-1^, to match the light intensity during natural crepuscular sunset.

To illuminate the fly and enable measurement of its wingbeat amplitude, a custom infra-red diffused backlight of 49 LEDs (Vishay, 2 in x 2 in, 880 nm) was positioned below the sphere. To capture the wingbeats, a high-speed camera (FL3-U3-13Y3M-C, Point Grey Research) with attached lens (Tamron macro lens, f/2, 60 mm) and an infrared pass-only filter (850 nm, Edmund Optics) was positioned above the sphere. The amplitudes of the fly’s left- and right-wingbeats were computed in real-time using custom software written in Microsoft Visual C++ using the machine vision OpenCV library at 200 Hz, allowing us to present visual stimuli both with and without yaw-turning closed-loop feedback.

Using a focusable dot infra-red laser (808 nm, maximum power 350 mW, Roithner LaserTechnik), we were able to deliver a heat punishment which was calibrated to raise the fly’s body temperature from ambient room temperature (25 °C) to 35 °C within 0.5s. Laser power to achieve this temperature increase was determined by inserting the top of a hypodermic needle thermocouple probe (HYP1-30-1/2-T-G-SMPW-M, Omega). An 850 nm long-pass dichroic beamsplitter (Edmund Optics) was placed in between the display sphere and the high-speed camera above it to direct the punishment laser light beam down on to the fly while still allowing for the camera above to detect the wing-beat image.

For optogenetic experiments, an additional 735-nm long-pass dichroic beamsplitter (Edmund Optics) was placed in the laser path - allowing red light stimulation (635 nm, Roithner LaserTechnik) to be pulsed at the fly head at 0.8 mW cm^-2^ for 0.5 sec.

### Conditioning protocols and learning performance

During the training phase, we presented either an upright-T or inverted-T shaped bright image over a dark background to naïve flies (not exposed to any conditioning previously). The Ts were displayed in the frontal visual field of the fly and were held fixed during presentation. The T-shape measured 40° vertically and horizontally, with the bars of the Ts being 14° wide. We randomized which image was used as CS^+^ (paired with heat), vs. the control CS^-^ condition (not paired with heat) between experiments. 7 training trials were performed for both delay and trace conditioning (5 s trace interval) experiments, each training trial lasting 60 s with a 30 s inter-trial-interval. Post-training, a single test of learning was performed for each conditioned fly. A 60 s gap was introduced between the end of the last training trial and start of the test sequence.

During the test phase, both upright-T and inverted-T shapes were presented simultaneously in closed-loop mode. The Ts were presented lateral to the fly and 180° apart, however which shape was displayed on the left vs. right of the fly was randomized between experiments. The performance index score (PI), a measure of learning was calculated as follows. If t_a_ was the time the fly spent orienting towards the CS^-^ image quadrant, and t_b_ was the time the fly spent orienting towards the CS^+^ image quadrant, PI was determined as (t_a_ - t_b_)/(t_a_ + t_b_). Therefore, PI > 0 indicated successful learning as the fly fixated more on CS^-^ than CS^+^, with PI < 0 indicating the opposite. PI = 0 indicated an equal probability of fixating on CS^+^ and CS^-^. Only experiments where the fly fixated (on either pattern quadrant) for at least 50 % of the total experiment time (10 out of 20 s) were considered in our analysis.

### Fly brain window surgery

A custom flyholder^67^ was used to immobilize the fly head (while allowing its wings to move freely) under an imaging device and expose its brain for optical recordings. The flyholder consisted of two parts – a 3D printed plastic frame, and a soft annealed stainless-steel shim (with a fly brain sized hole) folded and glued with epoxy to fit the contour of the frame^67^. After gluing the metal shim to the holder, charcoal primer and paint were applied to the bottom side of the shim to minimize reflections during wing-beat tracking.

Our fly mounting and surgery procedure is as described - cold-anesthetized flies (4 °C) were positioned ventral side up in a fly-sized divot machined in a custom-made brass block. The first pair of legs (T1) were cut at the first segment, and the middle and rear pairs (T2 and T3) were removed completely. The proboscis was gently pushed into the head capsule, and a small drop of UV-cured glue was applied to fix it in place. The fly was then flipped over, dorsal side up, and a small drop of UV-cured glue was applied in the gap between the head and thorax, thereby tilting the fly head slightly upwards. The flyholder was then positioned and glued to the head, following which, the holder and the fly were removed from cold-anesthesia and allowed to recover.

After recovery (determined by the fly exhibiting flight), the flyholder was filled with saline to fully cover the head. 1x saline containing 103 mM NaCl, 5 mM TES, 8 mM Trehalose, 10 mM Glucose, 26 mM NaHCO_3_, 1 mM NaH_2_PO_4_, 1.5 mM MgCl_2_, and 3mM CaCl_2_, with a pH = 7.3 was prepared and used. With the head covered in saline, the cuticle, air sacs and fat bodies were removed by hand dissection using fine forceps (Dumont 5SF, Fine Science Tools, or tip-size A, re-sharpened at Corte Instruments). An incision was made first along the posterior side of the head, then along both lateral edges, and finally along the anterior side to remove the cuticle.

### Two-photon *in-vivo* fluorescence imaging

The tethered flight behavior assay described above was coupled with a two-photon microscope with some modifications. First, a cylindrical acrylic display (4 in diameter, 3.5 in height, ∼330° of azimuth and ∼85° of elevation) with an adhesive rear-projection film was used for visual stimulus presentation. The display was tilted downwards (pitch angle of ∼20°) relative to the flyholder positioned in it. The projector used for visual stimulus presentation was identical to the tethered-flight assay except for a 447/60 bandpass filter (Chroma) that was placed in the light path to avoid any bleed through of visual stimulus light into the imaging recordings.

A collimated fiber-coupled 850 nm IR LED (M850F2, M28L01, F240SMA-850, Thorlabs), placed directly under the flyholder and behind the fly, was used to illuminate it for accurate positioning, the image of which was captured using a high-speed IR sensitive camera (FL3-U3-13Y3M-C, Point Grey Research). The IR laser punishment delivery apparatus (808 nm, maximum power 350 mW, Roithner LaserTechnik) was identical to the tethered flight assay and delivered from underneath the fly.

Our imaging experiments were performed using a Bruker Ultima Investigator multiphoton microscope with a Nikon 40x NIR Apo objective water-immersion lens (0.8 N.A., pixel size 0.27 x 0.27 μm, FOV 69.1 x 69.1 μm). GaAsP photomultiplier tubes (H10770, Hamamatsu) band-passed with either et525/70m-2p or et595/50m-2p emission filters (Chroma) and a t565lpxr dichroic beam-splitter (Chroma) were used for simultaneous acquisition. A 920 nm mode-locked Ti:Sapphire laser (Mai Tai, Spectra Physics) provided a maximum power of 15 mW on the sample. Resonant scanning galvos and a high-speed Z-piezo (Bruker, max range, 400 μm) were used for imaging a 6-plane volume of the EB at a rate of ∼6 Hz (image resolution of 256x256 px, 5 μm spacing between imaging planes).

We performed two broad classes of *in vivo* imaging experiments. First, ratiometric calcium imaging which consisted of multi-plane 3D volumes of calcium-dependent GCaMP and calcium-independent tdTomato fluorescence from the same neural populations, simultaneously captured with dual GaAsP PMTs. Second, Glutamate- and GABA-sensing fluorescence reporters (iGluSnFR and iGABASnFR2 respectively), wherein 3D volumes of the detectable fluorescence were recorded using the green-sensitive GaAsP PMT.

### Fluorescence quantification

We first generated a maximum intensity projection (MIP) sequence for each pixel (256x256 px) in the 3D volume for GCaMP/iGluSnFR/iGABASnFR2 and/or tdTomato channels. Each MIP sequence was filtered using a 3x3 median spatial filter followed by 2D gaussian smoothing with a square gaussian kernel standard deviation of 1, to smooth acquisition noise.

Next, for ratiometric calcium imaging experiments, the tdTomato channel sequence was then spatially aligned using a fast-normalized cross-correlation template matching algorithm based on the first MIP frame. The same transformations were then applied to the MIP sequence of the GCaMP channel to ensure pixel-pixel correspondence between frames of the two sequences. In case of SnFR reporter imaging experiments, the first MIP frame was used for the alignment.

We then applied (to the filtered raw data) a Principal Component Analysis (PCA) to decorrelate the data and uncover unknown, independent components of the observed features across time. We assume that the most discriminative information is captured by the largest variance in the feature space, an assumption likely true since the direction of the largest variance encodes the most information. We empirically determine the number of eigenvectors to consider and reduce the dimensionality of the raw data across all experiments to be 12, based on computing the total variance of the data represented by each ordered principal component.

The final step involved creating a binary mask of labeled neurons, segmenting foreground (neural) pixels from background. For ratiometric calcium imaging experiments, a mask was created by applying adaptive thresholding to each frame of the tdTomato channel. The calcium transient was then measured as a ratio, of average green calcium-dependent fluorescence (GCaMP) over red calcium-independent fluorescence (tdTomato) within the masked region. For SnFR reporter imaging experiments, an average intensity projection image of the MIP sequence was used to create a neural mask and determine average SnFR reporter transients within the masked region.

### Exponential model fitting

GCaMP/iGluSnFR/iGABASnFR2 change in fluorescence intensity levels were then determined by first applying a Savitzky-Golay filtering (order of 3, frame length of 7)^68^ to the raw intensity time-series data, then computing the difference of maximum F values from a baseline level: dF/F = (F - F_baseline)/F_baseline. Baseline levels were computed by averaging the lowest 25 % intensity frames during the baseline period (5 s) of the first trial before CS presentation.

Separate exponential curves were fit to segments of the dF/F data to determine decay/growth rates after termination of CS. The exponential decay/growth constant (tau), defined as the amount of time the activity signal would take to decay/grow by a factor of 1/e (approximately 36.8%), was used as a measure of signal decay/growth during learning.

### GCaMP frequency analysis

Frequency analyses were performed on the reduced dimensionality GCaMP data of ER2/4M neurons, as reported previously^64^, to quantify any increased oscillatory activity and signal persistence in those neurons during the conditioning process. Underlying frequencies were extracted by computing a discrete Fourier transform using the fast Fourier transform (FFT) algorithm. First, a baseline frequency spectrum of calcium activity prior to conditioning was computed using pre-CS (5 s) data (of the first trial) from for all experiments. Next, raw reduced dimensionality data were low pass filtered to exclude frequencies greater than 75 % of the Nyquist rate, which in our experiments was ∼3 Hz. This detectable frequency range is well within that of GCaMP6f intermediate kinetics which offers a temporal resolution (median ± s.e.m.) of 26 ± 2 ms (1 action-potential dF/F half-rise time) and 140 ± 20 ms (1 action-potential dF/F half-decay time^65^). The frequencies with maximum relative power above baseline were then obtained for different stages of the conditioning process.

### GluSnFR peak detection analysis

Local maxima peaks, defined as points where the value is higher than all nearby points, indicating a rise-then-fall pattern, were determined for glutamate-sensing fluorescence recordings. Minimum peak height and prominence thresholds of 5-standard-deviations of the corresponding baseline dF/F level were used to detect significant peaks in fluorescence. Peak prominence measured how much a peak stood out from the surrounding signal, defined as the vertical distance from the peak to the lowest contour line that encircled it and no higher peaks.

### Immunostaining

The following procedure was used for immunostaining whole-mount brains. First, brains dissected from 3-7 day old of female flies were fixed with 4 % paraformaldehyde for 2 hours at room temperature and followed by extensive washes with washing buffer (0.3 % PBT) for 4 x 20 min. After washes, brains were incubated in blocking buffer (5 % normal goat serum) for 1.5 hours. Next, the brains were incubated in primary and secondary antibodies at 4 °C for 48 hours each, with extensive washes (0.3 % PBT, 4 X 20 min) between steps. Primary antibodies used here included Rabbit anti-GFP (1:1000, Invitrogen) and mouse anti-nc82 (1:30, Developmental Studies Hybridoma Bank). Secondary antibodies used here were AF488 goat α-rabbit (1:400, Invitrogen), and AF568 goat α-mouse (1:400, Invitrogen). The brains were then mounted on microscope slides with Vectashield antifade mounting medium for fluorescence (Vector Laboratories).

### Confocal microscopy

Confocal laser scanning micrographs of immunostained whole-mount brains were obtained using a Zeiss LSM880 confocal microscope with a 20X (1.0 N.A.) water immersion objective. Spacing between individual planes was 1 μm and the pixel resolution was set to 1024 x 1024. At least three brains were immunostained for each genotype presented.

### Fly connectome analysis

The total connection strength between presynaptic cell type *A* and postsynaptic cell type *B* was calculated as follows. For each pair of neurons *i* ∈ *A* and *j* ∈ *B* , the signed connection strength *C_ij_* was computed as the product of the number of synapses *S_ij_* between neuron *i* and neuron *j* , and a sign *σ_ij_* determined by the neurotransmitter identity of the presynaptic neuron *i*,

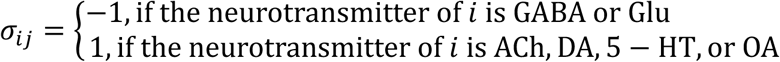

The total connectivity strength from cell type *A* to cell type *B* is then given by aggregating these pairwise values across all neuron pairs, *C_AB_* = ∑*_i_*_∈*A*_ ∑*_j_*_∈*B*_ *C_ij_*.

### Statistics and reproducibility

We performed all post-acquisition analyses using Matlab R2024a. Summarized data were represented either as means (with s.e.m) or boxplots, with center (median), edges (inter-quartile range), and whiskers (1.5x inter-quartile range).

When comparing more than two groups, Kruskall-Wallis and post-hoc unpaired two-sided Mann–Whitney U tests with bonferroni corrected multiple comparisons were used. Two group comparisons were performed using unpaired two-sided Mann Whitney U tests.

Behavioral and neurophysiological data with a two-factor structure (across trials and conditioning variants) were analyzed using two-factor aligned-rank-transform analysis of variance (ART-ANOVA)^69^. Single-factor comparisons (across trials for same conditioning paradigm) were performed using Friedman’s repeated measures ANOVA. Sample size estimates based on medium to large effect sizes (Cohen’s d), were calculated a priori, during the experimental design stage. Final sample sizes ensured a test power of at least 0.8. Experiments were performed non-blinded. Single fly metrics are overlaid in all figures as scatters. Immunostaining experiments were performed on at least three brains per genotype presented (see methods, immunostaining).

## Data availability

Datasets generated as part of this study will be made available from the corresponding author on reasonable request.

## Code availability

All source codes for the different assays used and analysis routines will be made available for download from the public repository at https://github.com/dgrover/flyWM.

## Acknowledgements

We thank G Turner for *UAS-iGABASnFR2*; NC Spitzer, RJ Greenspan, and members of the Grover laboratory for discussions and comments on the manuscript; J-Y Chen, S Godavarthi and W Choi for assistance with experiments; M Yao, TO Sharpee and K Asahina for discussions. This work was funded by Air Force Office of Scientific Research FA9550-23-1-0024 and FA9550-25-1-0299, Kavli Institute for Brain and Mind Innovative Research Grant 2021-1779 to DG.

## Author contributions

MLH performed the experiments with early support from JX. All authors were involved with study design and data analysis. DG supervised the study and MLH and DG authored the manuscript.

## Extended data figure legends

**Extended Data Figure 1. Sustained neural stability during trace conditioning requires feedforward GABAergic inhibition and co-transmission of glutamate from ER2/4m neurons. (a)** Ratiometric imaging of *EB1-Gal4>>UAS-GCaMP6f*.*myr-tdTomato* (left), *EB1-Gal4>>UAS-GCaMP6f*.*myr-tdTomato,UAS-Gad1^RNAi^*(center), and *EB1-Gal4>>UAS-GCaMP6f*.*myr-tdTomato,UAS-dVglut^RNAi^*(right) females during trace conditioning. Shown, dF_ratio_/F_ratio_ activity for trials 1 thru 7. Single-term exponential curve fits (red) through dF_ratio_/F_ratio_ activity starting at CS offset (see methods, fluorescence quantification). **(b)** dF_ratio_/F_ratio_ magnitude (y-intercept) of the exponential fit of ER2/4m calcium activity for *EB1-Gal4>>UAS-GCaMP6f*.*myr-tdTomato* (black, n = 12 flies), *EB1-Gal4>>UAS-GCaMP6f*.*myr-tdTomato,UAS-Gad1^RNAi^*(blue, n = 10 flies), and *EB1-Gal4>>UAS-GCaMP6f*.*myr-tdTomato,UAS-dVglut^RNAi^*(red, n = 11 flies) at (left) CS offset (start of trace interval, t = 15 s) and (right) US onset (end of trace interval, t = 20 s). **(c)** ER2/4m decay constants (tau) of exponential fit of dF_ratio_/F_ratio_ calcium activity for *EB1-Gal4>>UAS-GCaMP6f*.*myr-tdTomato* (black), *EB1-Gal4>>UAS-GCaMP6f*.*myr-tdTomato,UAS-Gad1^RNAi^*(blue), and *EB1-Gal4>>UAS-GCaMP6f*.*myr-tdTomato,UAS-dVglut^RNAi^*(red). Tau is defined as the amount of time the activity signal would take to decay/grow by a factor of 1/e (see methods, exponential model fitting). **(d)** Frequency of ER2/4m calcium activity for trace conditioning, during the trace interval, for *EB1-Gal4>>UAS-GCaMP6f*.*myr-tdTomato* (black), *EB1-Gal4>>UAS-GCaMP6f*.*myr-tdTomato,UAS-Gad1^RNAi^* (blue), and *EB1-Gal4>>UAS-GCaMP6f*.*myr-tdTomato,UAS-dVglut^RNAi^* (red) flies (see methods, GCaMP frequency analysis). Boxplot center (median), edges (IQR), whiskers (1.5x IQR). Scatters represent single-fly metrics. Groups compared using two-factor ART-ANOVA. ** indicates p-value < 0.01.

**Extended Data Figure 2. Feedforward GABAergic inhibition and co-transmission of glutamate from ER2/4m neurons are not required for delay conditioning. (a)** Ratiometric imaging of *EB1-Gal4>>UAS-GCaMP6f*.*myr-tdTomato* (left), *EB1-Gal4>>UAS-GCaMP6f*.*myr-tdTomato,UAS-Gad1^RNAi^*(center), and *EB1-Gal4>>UAS-GCaMP6f*.*myr-tdTomato,UAS-dVglut^RNAi^* (right) females during delay conditioning. Shown, dF_ratio_/F_ratio_ activity for trials 1 thru 7. Single-term exponential curve fits (red) through dF_ratio_/F_ratio_ activity starting at CS offset (see methods, fluorescence quantification). **(b)** dF_ratio_/F_ratio_ magnitude (y-intercept) of the exponential fit of ER2/4m calcium activity for *EB1-Gal4>>UAS-GCaMP6f*.*myr-tdTomato* (black, n = 10 flies), *EB1-Gal4>>UAS-GCaMP6f*.*myr-tdTomato,UAS-Gad1^RNAi^*(blue, n = 10 flies), and *EB1-Gal4>>UAS-GCaMP6f*.*myr-tdTomato,UAS-dVglut^RNAi^*(red, n = 10 flies) during the matched trace interval, at (left) CS offset (t = 15 s) and (right) 5 s after CS offset (t = 20 s). **(c)** ER2/4m decay constants (tau) of exponential fit of dF_ratio_/F_ratio_ calcium activity for *EB1-Gal4>>UAS-GCaMP6f*.*myr-tdTomato* (black), *EB1-Gal4>>UAS-GCaMP6f*.*myr-tdTomato,UAS-Gad1^RNAi^*(blue), and *EB1-Gal4>>UAS-GCaMP6f*.*myr-tdTomato,UAS-dVglut^RNAi^*(red). Tau is defined as the amount of time the activity signal would take to decay/grow by a factor of 1/e (see methods, exponential model fitting). **(d)** Frequency of ER2/4m calcium activity for delay conditioning, during the matched trace interval, for *EB1-Gal4>>UAS-GCaMP6f*.*myr-tdTomato* (black), *EB1-Gal4>>UAS-GCaMP6f*.*myr-tdTomato,UAS-Gad1^RNAi^*(blue), and *EB1-Gal4>>UAS-GCaMP6f*.*myr-tdTomato,UAS-dVglut^RNAi^*(red) flies (see methods, GCaMP frequency analysis). Boxplot center (median), edges (IQR), whiskers (1.5x IQR). Scatters represent single-fly metrics. Groups compared using two-factor ART-ANOVA. ** indicates p-value < 0.01.

**Extended Data Figure 3. Sustained neural stability during trace conditioning requires GABAergic inhibition of ER3/4d neurons and nitric oxide synthase-mediated signaling from these neurons. (a)** Ratiometric imaging of *c232-Gal4>>UAS-GCaMP6f*.*myr-tdTomato* (left), *c232-Gal4>>UAS-GCaMP6f*.*myr-tdTomato,UAS-Rdl^RNAi^* (center), and *c232-Gal4>>UAS-GCaMP6f*.*myr-tdTomato,UAS-dNOS^RNAi^*(right) females during trace conditioning. Shown, dF_ratio_/F_ratio_ activity for trials 1 thru 7. Single-term exponential curve fits (red) through dF_ratio_/F_ratio_ activity starting at CS offset (see methods, fluorescence quantification). **(b)** dF_ratio_/F_ratio_ magnitude (y-intercept) of the exponential fit of ER3/4d calcium activity for *c232-Gal4>>UAS-GCaMP6f*.*myr-tdTomato* (black, n = 13 flies), *c232-Gal4>>UAS-GCaMP6f*.*myr-tdTomato,UAS-Rdl^RNAi^*(blue, n = 9 flies), and *c232-Gal4>>UAS-GCaMP6f*.*myr-tdTomato,UAS-dNOS^RNAi^*(red, n = 11 flies) at (left) CS offset (start of trace interval, t = 15 s) and (right) US onset (end of trace interval, t = 20 s). **(c)** ER3/4d growth constants (tau) of exponential fit of dF_ratio_/F_ratio_ calcium activity for *c232-Gal4>>UAS-GCaMP6f*.*myr-tdTomato* (black), *c232-Gal4>>UAS-GCaMP6f*.*myr-tdTomato,UAS-Rdl^RNAi^*(blue), and *c232-Gal4>>UAS-GCaMP6f*.*myr-tdTomato,UAS-dNOS^RNAi^*(red). Tau is defined as the amount of time the activity signal would take to decay/grow by a factor of 1/e (see methods, exponential model fitting). Boxplot center (median), edges (IQR), whiskers (1.5x IQR). Scatters represent single-fly metrics. Groups compared using two-factor ART-ANOVA. ** indicates p-value < 0.01.

**Extended Data Figure 4. GABAergic inhibition of ER3/4d neurons and nitric oxide synthase-mediated signaling from these neurons is not required for delay conditioning. (a)** Ratiometric imaging of *c232-Gal4>>UAS-GCaMP6f*.*myr-tdTomato* (left), *c232-Gal4>>UAS-GCaMP6f*.*myr-tdTomato,UAS-Rdl^RNAi^*(center), and *c232-Gal4>>UAS-GCaMP6f*.*myr-tdTomato,UAS-dNOS^RNAi^* (right) females during delay conditioning. Shown, dF_ratio_/F_ratio_ activity for trials 1 thru 7. Single-term exponential curve fits (red) through dF_ratio_/F_ratio_ activity starting at CS offset (see methods, fluorescence quantification). **(b)** dF_ratio_/F_ratio_ magnitude (y-intercept) of the exponential fit of ER3/4d calcium activity for *c232-Gal4>>UAS-GCaMP6f*.*myr-tdTomato* (black, n = 12 flies), *c232-Gal4>>UAS-GCaMP6f*.*myr-tdTomato,UAS-Rdl^RNAi^* (blue, n = 7 flies), and *c232-Gal4>>UAS-GCaMP6f*.*myr-tdTomato,UAS-dNOS^RNAi^*(red, n = 7 flies) during the matched trace interval, at (left) CS offset (t = 15 s) and (right) 5 s after CS offset (t = 20 s). **(c)** ER3/4d decay constants (tau) of exponential fit of dF_ratio_/F_ratio_ calcium activity for *c232-Gal4>>UAS-GCaMP6f*.*myr-tdTomato* (black), *c232-Gal4>>UAS-GCaMP6f*.*myr-tdTomato,UAS-Rdl^RNAi^*(blue), and *c232-Gal4>>UAS-GCaMP6f*.*myr-tdTomato,UAS-dNOS^RNAi^*(red). Tau is defined as the amount of time the activity signal would take to decay/grow by a factor of 1/e (see methods, exponential model fitting). Boxplot center (median), edges (IQR), whiskers (1.5x IQR). Scatters represent single-fly metrics. Groups compared using two-factor ART-ANOVA. ** indicates p-value < 0.01.

**Extended Data Figure 5. Glutamate signaling shift prior to US onset observed selectively in trace conditioning. (a)** *In vivo* fluorescent glutamate indicator imaging from ER2/4m neurons, *EB1-Gal4>>UAS-iGluSnFR* female during delay conditioning (left) and trace conditioning (right). Shown, dF/F activity for trials 1 thru 7 with local maxima peaks (red circles) of dF/F activity (see methods, fluorescence quantification, and GluSnFR peak detection analysis). **(b)** Peak dF/F activity during CS (light red), and post-US (dark red) of delay conditioning (left, n = 7 flies); and during CS (light red), during TI (red), and post-US (dark red) of trace conditioning (right, n = 10 flies). **(c)** Timing of the highest local maxima peak of dF/F activity over trials during delay conditioning (left), and trace conditioning (right). In (b) and (c), quadratic (second degree) polynomial curve fits through median dF/F activity. Boxplot center (median), edges (IQR), whiskers (1.5x IQR). Scatters represent single-fly metrics. Groups compared using two-factor ART-ANOVA. ** indicates p-value < 0.01.

**Extended Data Figure 6. Sustained GABAergic signaling during trace conditioning is supported by glutamatergic co-transmission. (a)** *In vivo* fluorescent GABA indicator imaging from ER2/4m neurons, *EB1-Gal4>>UAS-iGluSnFR2* (left), and *EB1-Gal4>>UAS-iGABASnFR2,UAS-dVglut^RNAi^*(right) females, during trace conditioning. Shown, dF/F activity for trials 1 thru 7. Single-term exponential curve fits (red) through dF/F activity starting at CS offset (see methods, fluorescence quantification). **(b)** dF/F magnitude (y-intercept) of the exponential fit of ER2/4m GABAergic activity for *EB1-Gal4>>UAS-iGABASnFR2* (blue, n = 12 flies), and *EB1-Gal4>>UAS-iGABASnFR2,UAS-dVglut^RNAi^*(purple, n = 8 flies) at (left) CS offset (start of trace interval, t = 15 s) and (right) US onset (end of trace interval, t = 20 s). **(c)** ER2/4m decay constants (tau) of exponential fit of dF/F GABAergic activity for *EB1-Gal4>>UAS-iGABASnFR2* (blue), and *EB1-Gal4>>UAS-iGABASnFR2,UAS-dVglut^RNAi^* (purple). **(d)** dF/F magnitude (y-intercept) of the exponential fit of ER2/4m GABAergic activity for *EB1-Gal4>>UAS-iGABASnFR2* (blue, n = 12 flies) alongside dF_ratio_/F_ratio_ magnitude (y-intercept) of the exponential fit of ER2/4m calcium activity for *EB1-Gal4>>UAS-GCaMP6f*.*myr-tdTomato* (black, n = 12 flies) at (left) CS offset (start of trace interval, t = 15 s) and (right) US onset (end of trace interval, t = 20 s). **(e)** ER2/4m decay constants (tau) of exponential fit of dF/F GABAergic activity for *EB1-Gal4>>UAS-iGABASnFR2* (blue) alongside ER2/4m decay constants (tau) of exponential fit of dF_ratio_/F_ratio_ calcium activity for *EB1-Gal4>>UAS-GCaMP6f*.*myr-tdTomato* (black). Boxplot center (median), edges (IQR), whiskers (1.5x IQR). Scatters represent single-fly metrics. Groups compared using two-factor ART-ANOVA. ** indicates p-value < 0.01.

**Extended Data Figure 7. Sustained GABAergic signaling beyond CS offset not required for delay conditioning. (a)** *In vivo* fluorescent GABA indicator imaging from ER2/4m neurons, *EB1-Gal4>>UAS-iGluSnFR2* (left), and *EB1-Gal4>>UAS-iGABASnFR2,UAS-dVglut^RNAi^* (right) females, during delay conditioning. Shown, dF/F activity for trials 1 thru 7. Single-term exponential curve fits (red) through dF/F activity starting at CS offset (see methods, fluorescence quantification). **(b)** dF/F magnitude (y-intercept) of the exponential fit of ER2/4m GABAergic activity for *EB1-Gal4>>UAS-iGABASnFR2* (blue, n = 8 flies), and *EB1-Gal4>>UAS-iGABASnFR2,UAS-dVglut^RNAi^*(purple, n = 9 flies) during the matched trace interval, at (left) CS offset (t = 15 s) and (right) 5 s after CS offset (t = 20 s). **(c)** ER2/4m decay constants (tau) of exponential fit of dF/F GABAergic activity for *EB1-Gal4>>UAS-iGABASnFR2* (blue), and *EB1-Gal4>>UAS-iGABASnFR2,UAS-dVglut^RNAi^* (purple). **(d)** dF/F magnitude (y-intercept) of the exponential fit of ER2/4m GABAergic activity for *EB1-Gal4>>UAS-iGABASnFR2* (blue, n = 8 flies) alongside dF_ratio_/F_ratio_ magnitude (y-intercept) of the exponential fit of ER2/4m calcium activity for *EB1-Gal4>>UAS-GCaMP6f*.*myr-tdTomato* (black, n = 10 flies) during the matched trace interval, at (left) CS offset (start of trace interval, t = 15 s) and (right) US onset (end of trace interval, t = 20 s). **(e)** ER2/4m decay constants (tau) of exponential fit of dF/F GABAergic activity for *EB1-Gal4>>UAS-iGABASnFR2* (blue) alongside ER2/4m decay constants (tau) of exponential fit of dF_ratio_/F_ratio_ calcium activity for *EB1-Gal4>>UAS-GCaMP6f*.*myr-tdTomato* (black). Boxplot center (median), edges (IQR), whiskers (1.5x IQR). Scatters represent single-fly metrics. Groups compared using two-factor ART-ANOVA. ** indicates p-value < 0.01.

